# Enrichment of de novo mutations in non SNP sites in autism spectrum disorders and an empirical test of the neutral DNA model

**DOI:** 10.1101/231944

**Authors:** Ye Zhang, Shi Huang

**Affiliations:** Center for Medical Genetics, School of Life Sciences, Xiangya Medical School, Central South University, Changsha, Hunan 410078, People’s Republic of China

**Keywords:** autism, de novo mutations, parity rule, AT content, neutral theory, infinite site, maximum genetic diversity hypothesis

## Abstract

The genetic basis of autism spectrum disorders (ASD) remains better understood and might concern only a small fraction of the genome if the neutral theory were true. We here analyzed published de novo mutations (DNMs) in ASD and controls. We found that DNMs in normal subjects occurred at positions bearing SNPs at least 3.45 fold more frequent than expected from the neutral theory, whereas DNMs in ASD were less frequent relative to those in controls, especially so for common SNPs with minor allele frequency >0.01. Among sites bearing both SNPs and DNMs, DNMs in controls occurred significantly more frequent than DNMs in ASD at reference allele sites bearing C or G nucleotides, indicating depletion of ASD associated DNMs in known regions of hypermutability or less functional constraints such as CpG sites. We also analyzed the nucleotide compositions of DNMs and the parity (1:1 ratio) of pyrimidines and purines. We found that DNMs in ASD showed overall lower AT content than that in controls. Parity violations and AT bias in DNMs occurred at expected frequency based on chance in both ASD and controls. These results show enrichment of DNMs at positions bearing SNP sites and C or G sites in normal subjects and less so in ASD, which is not expected from the neutral model, and indicate that DNMs are on average more deleterious in ASD than in controls.

## Introduction

Autism spectrum disorders (ASD) is a common disease today with a prevalence of 14.6 per 1,000 (one in 68) children aged 8 years in the United States at 2012 ^1^. It is four times more common in males than in females ^2, 3^. Twin and family studies show that siblings of children with ASD are at a significant higher risk for autism than the general population ^4–6^. ASD remains poorly understood but may have a strong genetic component with a heritability of 40–80% ^7–10^. ASD are genetically highly heterogeneous, with no single gene accounting for more than 1% of cases ^11^.

Recent work has shown a substantial contribution of de novo variations ^12–15^. Probands of ASD, relative to unaffected siblings, have been found to more likely carry multiple coding and noncoding DNMs in different genes, which are enriched for expression in striatal neurons ^16^. Genome-wide association studies have revealed few replicable common polymorphisms associated with ASD ^17–20^. Common genetic variants are individually of little effect but acting additively may be a major source of risk for autism ^21^. Assortative mating may play a role in bringing about enrichment of ASD alleles in an affected child ^22^. Consistent with the notion of collective and additive effects of common variants, recent studies indicate a role for genome wide minor allele content (MAC) of an individual in a variety of complex traits and diseases ^23–26^. The more the number of minor alleles of common SNPs in an individual, the higher the risk in general for many complex diseases such as type 2 diabetes, schizophrenia, and Parkinson’s disease ^23–28^. Such findings indicate an optimum level of genetic variations that an individual can tolerate ^29, 30^.

Nucleotide positions of common SNPs found in normal populations such as in the 1000 genomes cohort are presumably less conserved than those regions of genome that are deficient in SNPs. The neutral theory has served as the theoretical foundation for the inference that most of the human genome (~90%) are freely changeable or selectively neutral ^31^. However, it remains to be empirically verified whether the SNP depleted regions of the genome can freely tolerate DNMs as expected from such inference. The neutral theory is widely thought to have failed to explain the genetic diversity riddle and other major evolutionary phenomenon ^32–36^. While a small fraction of DNMs in ASD have been found to be enriched in deleterious mutations relative to those in normal subjects ^16^, it remains unknown whether the rest or most of DNMs are also deleterious or occur more often in the genomic regions deficient in SNPs or are overall different from DNMs in controls.

In this study, we investigated whether DNMs in ASD are more enriched in the SNP deficient sites relative to those in normal subjects and whether DNMs in normal subjects may show preference for positions bearing the common SNPs. We also studied nucleotide composition patterns in DNMs in ASDs in terms of AT content (human genome is known to be ~58% AT) and Chargaff’s Parity Rule 1 and 2 (the 1:1 ratio of pyrimidines and purines) ^37, 38^. We found that DNMs in ASDs were more enriched in positions deficient in common SNPs relative to those in controls and that DNMs in normal individuals occurred more often in the common SNP sites. DNMs in ASD showed lower AT content but normal parity patterns. The results do not support the inference of 90% non-constrained genome as inferred from the neutral model.

## Results

### DNMs were enriched in sites of common SNPs and less so in ASD

We made use of the DNM database NPdenov ^39^ to study the genomic mutation patterns in ASD (http://www.wzgenomics.cn/NPdenovo/). We focused on SNVs and studied 50281 DNMs in control subjects and 28376 DNMs in ASD that were discovered by whole genome sequencing. We matched the positions of these DNMs with those bearing SNPs detected in the 1000 genomes project (1KGP) phase 3 dataset ^40^. The fraction of DNM sites matching the SNP sites of 1KGP (total SNPs numbers 81,377,202) in ASD was found lower than that in the control subjects (2056/28376 or 0.073 vs 4457/50280 or 0.089, P < 0.001, chi square test, Table 1). For SNPs with minor allele frequency (MAF) >0.01 in the 1KGP (numbers 12,200,686), the matches were only 108/28376 or 0.0038 for DNM in ASD versus 354/50280 or 0.0070 for DNM in controls (P < 0.001, chi square test). Also in the case of ASD, the fraction of all SNPs matched with DNMs was 2.8 times that of higher MAF (>0.01) SNPs matched with DNMs (0.000025 vs 0.0000089, P<0.01), whereas in the case of controls, the fraction of all SNPs matched with DNMs was only 1.9 times that of higher MAF (>0.01) SNPs matched with DNMs (0.000055 vs 0.000029, P<0.01, Table 1), indicating again that DNMs in ASD cases occurred more often in the rare SNP sites.

**Table 1.** Match of DNMs with common SNPs

| Fractions matched | ASD cases |  | Controls |  |
| --- | --- | --- | --- | --- |
|  | DNM | SNPs | DNM | SNPs |
| All SNPs | 0.0725 |  | 0.089*** |  |
| All DNMs |  | 0.000025 |  | 0.000055 |
| SNPs (MAF>0.01) | 0.0038 |  | 0.007*** |  |
| DNMs (MAF>0.01) |  | 0.0000089** |  | 0.000029** |

\*\*\*, P< 0.001, ASD cases vs controls, chi squared test; \*\*, P< 0.01, All SNPs vs SNPs with MAF>0.01.

These results show a preferential depletion of DNMs in positions bearing SNPs, especially for those bearing the common SNPs (MAF>0.01), in ASD relative to controls.

We analyzed 81.4 million SNPs in 1KGP representing 2.58% of the total number of bases in the GRCh37 hg19 genome (3.156 billion). If most sites in the genome are neutral and can accommodate mutations equally, one would expect ~2.58% of DNMs to overlap with SNPs. In fact, the infinite site model, which is compatible with the neutral framework and widely used to interpret observed polymorphisms, predicts this percentage to be much lower since the model means that new mutations should mostly occur at never before mutated sites. However, the observed percentage in normal subjects, 8.9% (Table 1), was 3.45 fold higher than the expected value, which was likely an underestimation. This could be accounted for only if just 29% of the genome can freely accommodate mutations.

There were 12.2 million SNPs with MAF>0.01 in 1KGP representing 0.39% of the genome. The observed percentage of DNMs matching SNPs with MAF>0.01 in normal subjects was 0.7% in normal subjects and 0.38% in ASD cases. So, the percentage in normal subjects was 1.81 fold higher than expected while the percentage in ASD cases was similar to or slightly lower than the expected value. These observations show that DNMs in normal subjects did not, whereas DNMs in ASD cases did to some extent, conform to inferences by the infinite site model and the neutral theory. So, most parts of the genome (71%) may not freely tolerate mutations that can survive as DNMs in healthy individuals. If de novo mutations do occur mostly on new sites as predicted by the infinite site model, they would produce a pattern similar to that in the ASD cases but unlike that in the normal subjects, and hence be associated with diseases.

There are hyper-mutable regions in the genome and CpG sites are known to have higher mutation rates ^41^. Such regions would be more tolerable to mutations or under less functional constraint and hence expected to be enriched with SNPs. If DNMs in ASD tend to cluster in functionally constrained sites, they should overlap less with those SNP sites bearing C or G. We therefore examined among all the DNMs matched with SNP sites the fraction of DNMs that have reference allele being C, G, A, or T nucleotide (Figure 1). The results showed that the fraction of DNMs that had C or G but not A or T reference alleles was significantly higher in controls than in ASD cases, indicating less enrichment of DNMs in ASD cases in hyper mutable sites. Also, hypermutability did explain a part of the match of DNMs with SNPs as reference alleles carrying C or G had higher fractions of match with SNP sites than those carrying A or T (Figure 1). The enrichment of DNMs in hypermutable regions further confirm that the non constrained regions in the genome may be much smaller than that expected from the inference based on the neutral framework.

**Figure 1.**
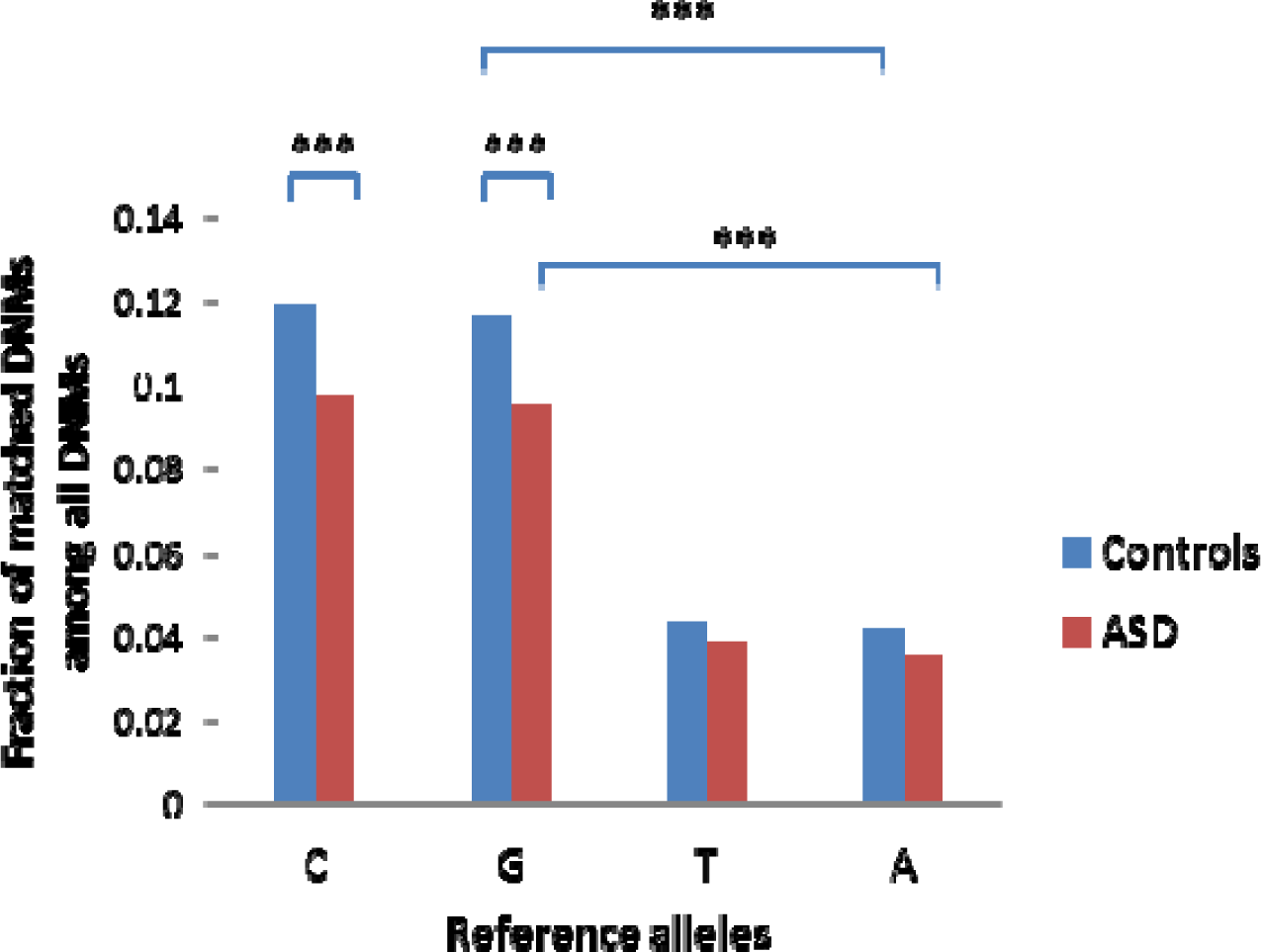
Fraction of DNMs with reference allele being C, G, A, or T nucleotide among all DNMs matched with SNP sites in 1KGP. ***, P < 0.001, chi-squared test, 2 tailed. The fractions of C or G were all significantly higher than that of A or T with P value shown for only some comparisons due to space constraints.

### DNMs in ASD were AT deficient but conformed to parity rules

The human genome shows unexplained AT-bias in base compositions (~58%). However, the overall AT content in DNMs in ASD cases showed less AT than controls (16017/28376 or 56.4% in ASD vs 28754/50281 or 57.2% AT in controls, P<0.05). To study AT content variation of DNMs among individuals, we examined 290 normal and 429 ASD individuals with 30-100 DNMs per individual. Among 429 ASD samples studied (Table 2), 7 (1.6%) showed AT bias (defined as higher than 58% AT, P<0.05) and 28 (6.5%) showed GC bias (defined as higher than 42% GC, P<0.05). In comparison, among 290 controls studied, two (0.69%) showed AT bias and 12 (4.1%) showed GC bias (P<0.05). The incidence of GC bias in ASD was higher than that in controls (P<0.05, chi squared test). The results showed that DNMs in ASD were significantly different from those in normal subject in being AT deficient.

**Table 2.** AT contents analyses.

|  | AT % | AT bias % | GC bias % |
| --- | --- | --- | --- |
| ASD | 56.4 | 1.6 | 6.5 |
| Controls | 57.2 | 0.69 | 4.1 |
| P value | < 0.05 | > 0.05 | < 0.05 |

DNMs are thought to occur randomly and expected to follow the 1:1 ratio of pyrimidines and purines (Chargaff’s Parity Rule 1 and 2). Random mutation process predicts that parity should hold for bases either targeted for (reference alleles) or resulting from mutations (alternative alleles). We confirmed this in our survey of 50281 DNMs in normal individuals as reported in the literature (25016 TC vs 25265 AG for reference alleles, and 25140 TC vs 25141 AG for alternative alleles). This pattern also held similarly in 28376 DNMs in ASD (14167 TC vs 14209 AG for reference alleles, and 14308 TC vs 14068 AG for alternative alleles). Therefore, one expects that mutations causing parity violations would qualify as non-random, in the same sense as a biased coin toss. We examined 290 normal and 429 ASD individuals with 30-100 DNMs per individual and found 4.8% (14/290) and 4.2 % (18/429) with parity violations (P<0.05), respectively. However, none of these were significant after adjustment for multiple testing. The rate of ~5% parity violations in the general population or ASD cases was consistent with the random chance of a nonrandom event as defined by P < 0.05. The results indicated that DNMs in ASD did not differ in conforming to parity rules from those in control subjects.

Of 18 ASD samples with parity violations, none showed AT bias and 4 showed GC bias (GC bias as defined by significantly greater than 42%, P<0.05). Of 14 normal samples with parity violations, none showed AT bias and one showed GC bias. Thus, of 32 samples with parity violations among 719 samples examined (290 normal and 429 ASD), none showed both parity violation and AT bias (incidence rate < 1/719 or 0.0014). Given the observed random rate of violating the parity rule and the AT bias pattern being 0.0445 and 0.0125, respectively, one would expect the random rate of violating both parity and AT bias pattern to be 0.00056, too low to be observed with the sample size studied here (719).

### Discussion

To better understand the genetics of ASD and the issue of neutral DNAs, we studied the genomic patterns of DNMs in ASD and normal subjects. Our results showed that relative to controls DNMs in ASD were more enriched in the conserved and less mutable regions of the genome that are deficient in common SNPs, which is consistent with previous findings of ASD probands carrying more DNMs with deleterious effects ^16^. This suggests that the conserved regions of the genome are under natural selection.

The neutral model explains the extremely low genetic diversity of humans, in particular non-Africans, by postulating bottlenecks in the past. This implies that the genetic diversity of non-Africans today would be much higher or similar to Africans if there was not bottleneck in the past. It also means that with more time to evolve the genetic diversity of non-Africans would be much greater in the future than it is now today. Based on the inference of 90% non-constrained genome under the neutral framework ^31^, one would expect most of the regions that are devoid of SNPs, which is greater than 90% based on the 1KGP data, to be free from natural selection. Under such inference, one would not necessarily expect the locations of most DNMs in ASD to be different from those in control subjects. In particular, the infinite site assumption of the neutral model is a prerequisite for most studies in the population genetics field and predicts that most DNMs should occur at new sites (have not had mutations before) and hence not to be enriched in SNP sites ^42^. In contrast to the inference of only 10% functional human genome based on the neutral theory ^31^, the maximum genetic diversity (MGD) hypothesis postulates that nearly all DNAs in humans are functional ^43, 44^. As mutations in functional DNAs are likely to be deleterious, the fraction of deleterious mutations among all new mutations would be similar to the fraction of the genome that is functional. Hence, the MGD theory predicts that most mutations and SNPs are deleterious, whereas the inference based on the neutral theory predicts only 10% of all new mutations to be deleterious. The MGD theory further postulates that genetic diversity must have an optimum upper limit and that genetic diversity in humans has reached saturation today. Therefore, the MGD hypothesis predicts that DNMs should be more enriched in positions bearing SNPs and that recurrent mutations should be common.

Our results here indicate that 71% of the genome may not be free to incur de novo mutations, which is much greater than the fraction of the genome (~10%) that is estimated to be functional by the neutral theory ^31^. The results therefore invalidate the neutral model and its infinite site assumption and support the MGD hypothesis. That SNP sites were preferentially hit by DNMs was consistent with the finding of saturated or maximum genetic diversity as observed in present day human populations ^45^. Sharing of common SNPs between different populations appear to be due to recurrent mutations rather than common ancestry. The neutral null hypothesis has been mostly tested previously by computational approaches based on uncertain assumptions that take certain sequences to be neutral for granted (repeats, viral sequences, and non-conserved regions) ^31^. Recent empirical studies comparing genetic diversities of patient and control populations have all contradicted the neutral model ^25, 27, 28, 30^. The results here provide additional empirical evidence not favoring the neutral hypothesis. The rapid advances in genomics in recent decades have made it practically possible for the first time to empirically test most of the uncertain assumptions in the field of population genetics and molecular evolution, and we expect more experimental tests to be performed in the near future. Reestablishing more realistic and certain assumptions will be key for the field to produce realistic conclusions that can actually find support from findings in other fields.

Because the collective effect of SNPs in numerous complex traits and diseases ^25, 27, 28, 30, 46^, even common SNPs are also mostly not neutral and only more neutral relative to the more conserved regions of the genome. Such observations therefore would further substantially reduce the fraction of neutral sites in the human genome.

Base mutations appear to have a bias in the direction of A:T and newly emerged low-frequency SNP alleles are typically A:T rich ^47, 48^. Interestingly, derived species appear to show more AT bias than ancestral species, and the direction of evolution appears to be going towards higher AT content ^38^. Our observation here of less AT content in DNMs of ASD indicates that AT content in DNMs of ASD was abnormal and against the trend of evolution. This may be related to the enigma of how AT content stays at a certain biased level not expected from random chance. Negative selection due to diseases such ASD may prevent AT content from increasing without limit. Our results is consistent with a previous study showing that ASD genes are AT rich ^49^. Thus, if mutations occurred in these AT rich ASD genes, there would be a high probability of mutating non-AT sites, leading to DNMs in these genes to be less rich AT.

Parity rules appear to apply in DNMs in both ASD and control subjects. Individuals with parity violations or AT bias exist in frequencies expected by random chance (~0.5%) in both ASD and controls. This confirms that DNMs occur mostly in a random fashion and obey probability rules. The event of both parity violation and AT bias in DNMs in an individual appears to be rare and future larger sample size studies are required to reveal whether such event may occur in reality.

## Materials and Methods

DNMs data from ASD and normal subjects were downloaded from NPdenovo database (http://www.wzgenomics.cn/NPdenovo/) ^39^. These DNMs were all found by whole genome sequencing analyses. For SNPs, we downloaded vcf files of the 1KGP phase 3 dataset ^40^. We used MAF data in the vcf file based on all ~2500 individuals in the 1KGP.

Data manipulations were done using custom scripts. Standard statistics methods were used including chi squared test (2 tailed) and Bonferroni correction for multiple testing and the statistics software Graphpad Prism6 was used for these analyses.

## Acknowledgments

We thank Jinchen Li for help with the NPdenovo database. This work was supported by the National Natural Science Foundation of China grant 81171880 and the National Basic Research Program of China grant 2011CB51001 (S.H.).

## Conflict of Interest Statements

The authors declare that they have no competing interests.

## Author Contributions

YZ and SH designed the analysis, analyzed data and wrote paper.

## References

1. Christensen DL, Baio J, Van Naarden Braun K, Bilder D, Charles J, Constantino JN et al. Prevalence and Characteristics of Autism Spectrum Disorder Among Children Aged 8 Years--Autism and Developmental Disabilities Monitoring Network, 11 Sites, United States, 2012. MMWR Surveill Summ 2016; 65(3): 1–23.

2. Lord C, Schopler E, Revicki D. Sex differences in autism. J Autism Dev Disord 1982; 12(4): 317–330.

3. Wing L, Wing JK. Early childhood autism: clinical, educational, and social aspects. 2d edn. Pergamon Press: Oxford; New York, 1976, xii, 342 p.pp.

4. Wood CL, Warnell F, Johnson M, Hames A, Pearce MS, McConachie H et al. Evidence for ASD recurrence rates and reproductive stoppage from large UK ASD research family databases. Autism Res 2015; 8(1): 73–81.

5. Bailey A, Le Couteur A, Gottesman I, Bolton P, Simonoff E, Yuzda E et al. Autism as a strongly genetic disorder: evidence from a British twin study. Psychol Med 1995; 25(1): 63–77.

6. Geschwind DH. Advances in autism. Annu Rev Med 2009; 60: 367–380.

7. Geschwind DH. Genetics of autism spectrum disorders. Trends Cogn Sci 2011; 15(9): 409–416.

8. Hallmayer J, Cleveland S, Torres A, Phillips J, Cohen B, Torigoe T et al. Genetic heritability and shared environmental factors among twin pairs with autism. Arch Gen Psychiatry 2011; 68(11): 1095–1102.

9. Robinson EB, Koenen KC, McCormick MC, Munir K, Hallett V, Happe F et al. A multivariate twin study of autistic traits in 12-year-olds: testing the fractionable autism triad hypothesis. Behav Genet 2012; 42(2): 245–255.

10. Gaugler T, Klei L, Sanders SJ, Bodea CA, Goldberg AP, Lee AB et al. Most genetic risk for autism resides with common variation. Nat Genet 2014; 46(8): 881–885.

11. Chahrour MH, Yu TW, Lim ET, Ataman B, Coulter ME, Hill RS et al. Whole-exome sequencing and homozygosity analysis implicate depolarization-regulated neuronal genes in autism. PLoS Genet 2012; 8(4): e1002635.

12. Iossifov I, O’Roak BJ, Sanders SJ, Ronemus M, Krumm N, Levy D et al. The contribution of de novo coding mutations to autism spectrum disorder. Nature 2014; 515(7526): 216–221.

13. Sanders SJ, Murtha MT, Gupta AR, Murdoch JD, Raubeson MJ, Willsey AJ et al. De novo mutations revealed by whole-exome sequencing are strongly associated with autism. Nature 2012; 485(7397): 237–241.

14. O’Roak BJ, Vives L, Girirajan S, Karakoc E, Krumm N, Coe BP et al. Sporadic autism exomes reveal a highly interconnected protein network of de novo mutations. Nature 2012; 485(7397): 246–250.

15. Neale BM, Kou Y, Liu L, Ma’ayan A, Samocha KE, Sabo A et al. Patterns and rates of exonic de novo mutations in autism spectrum disorders. Nature 2012; 485(7397): 242–245.

16. Turner TN, Coe BP, Dickel DE, Hoekzema K, Nelson BJ, Zody MC et al. Genomic Patterns of De Novo Mutation in Simplex Autism. Cell 2017; 171(3): 710–722 e712.

17. Devlin B, Melhem N, Roeder K. Do common variants play a role in risk for autism? Evidence and theoretical musings. Brain Res 2011; 1380: 78–84.

18. Pan Y, Chen J, Guo H, Ou J, Peng Y, Liu Q et al. Association of genetic variants of GRIN2B with autism. Sci Rep 2015; 5: 8296.

19. Xia K, Guo H, Hu Z, Xun G, Zuo L, Peng Y et al. Common genetic variants on 1p13.2 associate with risk of autism. Mol Psychiatry 2014; 19(11): 1212–1219.

20. Wang K, Zhang H, Ma D, Bucan M, Glessner JT, Abrahams BS et al. Common genetic variants on 5p14.1 associate with autism spectrum disorders. Nature 2009; 459(7246): 528–533.

21. Klei L, Sanders SJ, Murtha MT, Hus V, Lowe JK, Willsey AJ et al. Common genetic variants, acting additively, are a major source of risk for autism. Mol Autism 2012; 3(1): 9.

22. Zhu Z, Lu X, Yuan D, Huang S. Close genetic relationships between a spousal pair with autism-affected children and high minor allele content in cases in autism-associated SNPs. Genomics 2017; 109(1): 9–15.

23. Zhu Z, Man X, Xia M, Huang Y, Yuan D, Huang S. Collective effects of SNPs on transgenerational inheritance in Caenorhabditis elegans and budding yeast. Genomics 2015; 106(1): 23–29.

24. Zhu Z, Lu Q, Wang J, Huang S. Collective effects of common SNPs in foraging decisions in Caenorhabditis elegans and an integrative method of identification of candidate genes. Sci Rep 2015; 5: 16904.

25. Zhu Z, Yuan D, Luo D, Lu X, Huang S. Enrichment of Minor Alleles of Common SNPs and Improved Risk Prediction for Parkinson’s Disease. PLoS One 2015; 10(7): e0133421.

26. Yuan D, Zhu Z, Tan X, Liang J, Zeng C, Zhang J et al. Scoring the collective effects of SNPs: association of minor alleles with complex traits in model organisms. Sci China Life Sci 2014; 57(9): 876–888.

27. He P, Lei X, Yuan D, Zhu Z, Huang S. Accumulation of minor alleles and risk prediction in schizophrenia. Sci Rep 2017; 7(1): 11661.

28. Lei X, Huang S. Enrichment of minor allele of SNPs and genetic prediction of type 2 diabetes risk in British population. PLoS One 2017; 12(11): e0187644.

29. Hu T, Long M, Yuan D, Zhu Z, Huang Y, Huang S. The genetic equidistance result: misreading by the molecular clock and neutral theory and reinterpretation nearly half of a century later. Sci China Life Sci 2013; 56(3): 254–261.

30. Huang S. New thoughts on an old riddle: What determines genetic diversity within and between species? Genomics 2016; 108(1): 3–10.

31. Ponting CP, Hardison RC. What fraction of the human genome is functional? Genome Res 2011; 21(11): 1769–1776.

32. Hahn MW. Toward a selection theory of molecular evolution. Evolution 2008; 62(2): 255–265.

33. Kern AD, Hahn MW. The Neutral Theory in Light of Natural Selection. Mol Biol Evol 2018; 35(6): 1366–1371.

34. Kreitman M. The neutral theory is dead. Long live the neutral theory. Bioessays 1996; 18(8): 678–683; discussion 683.

35. Leffler EM, Bullaughey K, Matute DR, Meyer WK, Segurel L, Venkat A et al. Revisiting an old riddle: what determines genetic diversity levels within species? PLoS Biol 2012; 10(9): e1001388.

36. Ohta T, Gillespie JH. Development of Neutral and Nearly Neutral Theories. Theor Popul Biol 1996; 49(2): 128–142.

37. Rudner R, Karkas JD, Chargaff E. Separation of B. subtilis DNA into complementary strands. 3. Direct analysis. Proc Natl Acad Sci U S A 1968; 60(3): 921–922.

38. Li X, Scanlon MJ, Yu J. Evolutionary patterns of DNA base composition and correlation to polymorphisms in DNA repair systems. Nucleic Acids Res 2015; 43(7): 3614–3625.

39. Li J, Cai T, Jiang Y, Chen H, He X, Chen C et al. Genes with de novo mutations are shared by four neuropsychiatric disorders discovered from NPdenovo database. Mol Psychiatry 2016; 21(2): 290–297.

40. Auton A, Brooks LD, Durbin RM, Garrison EP, Kang HM, Korbel JO et al. A global reference for human genetic variation. Nature 2015; 526(7571): 68–74.

41. Nachman MW, Crowell SL. Estimate of the mutation rate per nucleotide in humans. Genetics 2000; 156(1): 297–304.

42. Kimura M, Crow JF. The Number of Alleles That Can Be Maintained in a Finite Population. Genetics 1964; 49: 725–738.

43. Huang S. Histone methylation and the initiation of cancer. CRC Press: New York, 2008.

44. Huang S. Inverse Relationship Between Genetic Diversity and Epigenetic Complexity. Available from Nature Precedings http://dx.doi.org/10.1038/npre.2009.1751.2, 2009.

45. Yuan D, Lei X, Gui Y, Zhu Z, Wang D, Yu J et al. Modern human origins: multiregional evolution of autosomes and East Asia origin of Y and mtDNA. bioRxiv 2017.

46. Lei X, Yuan D, Zhu Z, Huang S. Collective effects of common SNPs and risk prediction in lung cancer. Heredity (Edinb) 2018.

47. Hershberg R, Petrov DA. Evidence that mutation is universally biased towards AT in bacteria. PLoS Genet 2010; 6(9): e1001115.

48. Lynch M. Rate, molecular spectrum, and consequences of human mutation. Proc Natl Acad Sci U S A 2010; 107(3): 961–968.

49. Jabbari K, Nurnberg P. A genomic view on epilepsy and autism candidate genes. Genomics 2016; 108(1): 31–36.

